# A putative microcin amplifies Shiga toxin 2a production of *Escherichia coli* O157:H7

**DOI:** 10.1101/646182

**Authors:** Hillary M. Figler, Lingzi Xiaoli, Kakolie Banerjee, Maria Hoffmann, Kuan Yao, Edward G. Dudley

## Abstract

*Escherichia coli* O157:H7 is a foodborne pathogen, implicated in various multi-state outbreaks. It encodes Shiga toxin on a prophage, and Shiga toxin production is linked to phage induction. An *E. coli* strain, designated 0.1229, was identified that amplified Stx2a production when co-cultured with *E. coli* O157:H7 strain PA2. Growth of PA2 in 0.1229 cell-free supernatants had a similar effect, even when supernatants were heated to 100°C for 10 min, but not after treatment with Proteinase K. The secreted molecule was shown to use TolC for export and the TonB system for import. The genes sufficient for production of this molecule were localized to a 5.2 kb region of a 12.8 kb plasmid. This region was annotated, identifying hypothetical proteins, a predicted ABC transporter, and a cupin superfamily protein. These genes were identified and shown to be functional in two other *E. coli* strains, and bioinformatic analyses identified related gene clusters in similar and distinct bacterial species. These data collectively suggest *E. coli* 0.1229 and other *E. coli* produce a microcin that induces the SOS response in target bacteria. Besides adding to the limited number of microcins known to be produced by *E. coli*, this study provides an additional mechanism by which *stx2a* expression is increased in response to the gut microflora.

**Importance:** How the gut microflora influences the progression of bacterial infections is only beginning to be understood. Antibiotics are counter-indicated for *E. coli* O157:H7 infections, and therefore treatment options are limited. An increased understanding of how the gut microflora directs O157:H7 virulence gene expression may lead to additional treatment options. This work identified *E. coli* that enhance the production of Shiga toxin by O157:H7, through the secretion of a proposed microcin. This work demonstrates another mechanism by which non-O157 *E. coli* strains may increase Shiga toxin production, and adds to our understanding of microcins, a group of antimicrobials that are less well understood than colicins.

## Introduction

*E. coli* O157:H7 is a notorious member of the enterohemorrhagic *E. coli* (EHEC) pathotype, which causes hemolytic colitis and hemolytic uremic syndrome (HUS) through production of virulence factors including the locus of enterocyte effacement (LEE) and Shiga toxins (Stx) (1, 2). Stx is encoded on a lambdoid prophage (3). Induction of the prophage and subsequent upregulation of *stx* is tied to activation of the bacterial SOS response (4). Therefore, DNA damaging agents including certain antibiotics increase Stx synthesis, and are typically counter indicated during treatment (5). There are two Stx types, referred to as Stx1 and Stx2 (6). Stx1 is further divided into three subtypes, Stx1a, Stx1c and Stx1d (7). Stx2 also has multiple subtypes, designated Stx2a, Stx2b, Stx2c, Stx2d, Stx2e, Stx2f, Stx2g (7), Stx2h (8) and Stx2i (9). In general, infections caused by Stx1, and interestingly, even those with both Stx1 and Stx2 (such as strains EDL933 (10) and Sakai (11)) are associated with less severe disease symptoms than Stx2-only producing *E. coli* (12–14). Of the Stx2 subtypes, Stx2a is more commonly associated with clinical cases and instances of HUS (14–17). Indeed, the FAO and WHO considers STEC carrying *stx2a* to be of greatest concern (18).

Stx2a levels can be affected *in vitro* and *in vivo* when *E. coli* O157:H7 is cultured along with other bacteria. Indeed, it was found that *stx2a* expression is downregulated by various probiotic species (19, 20) or in a media conditioned with human microbiota (21). Conversely, non-pathogenic *E. coli* that are susceptible to infection by the *stx2a*-converting phage were reported to increase Stx2a levels (22, 23). This mechanism is O157:H7 strain-dependent (23), and requires expression of the *E. coli* BamA, which is the phage receptor (24, 25).

Production of Stx2a by O157:H7 can also increase in response to molecules secreted by other members of the gut microbiota (24, 26), such as bacteriocins and microcins. Bacteriocins are proteinaceous toxins produced by bacteria that inhibit the growth of closely related bacteria. For example, a colicin E9 (ColE9) producing strain amplified Stx2a when grown together with Sakai to higher levels than a colicin E3 (ColE3) producing strain (26). ColE9 is a DNase, while ColE3 has RNase activity, and this may explain the differences in SOS induction and Stx2a levels. In support of this, the addition of extracted DNase colicins to various *E. coli* O157:H7 strains increased Stx2a production, but not Stx1 (26). Additionally, microcin B17 (MccB17), a DNA gyrase inhibitor, was shown to amplify Stx2a production (24).

This work is part of a continuing effort to identify mechanisms by which members of the gut microbiome, especially *E. coli*, enhance Stx2a production of *E. coli* O157:H7. Of particular interest was the identification of secreted enhancer molecules, which were not investigated in our earlier studies. It was hypothesized that non-pathogenic *E. coli* strains could secrete additional colicins and microcins capable of increasing Stx2a production by O157:H7.

## Results

### 0.1229 amplifies Stx2a production in a cell independent manner

Human-associated *E. coli* isolates were tested for their ability to enhance Stx2a production in co-culture with O157:H7. Strain 0.1229 significantly increased Stx2a production of PA2, compared to PA2 alone (Fig. 1). C600 was included as a positive control, as it was previously shown to increase Stx2a production when co-cultured with O157:H7 (22, 23).

**Fig. 1:**
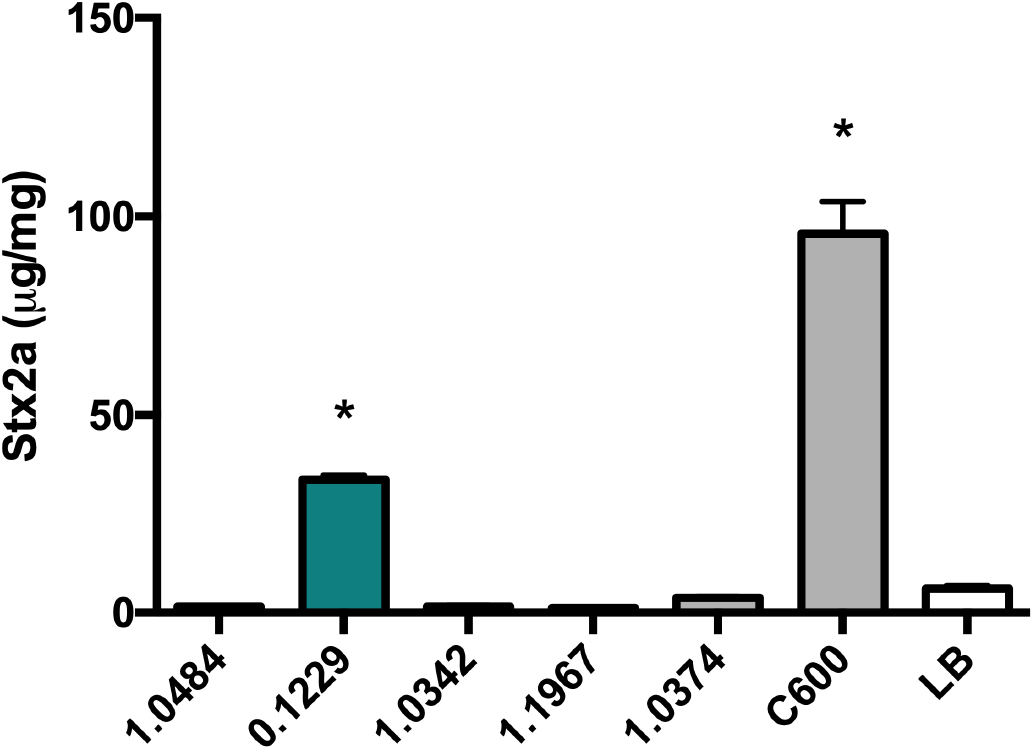
PA2 was grown with various *E. coli* strains and Stx2a levels were measured using the R-ELISA. LB refers to PA2 grown in mono-culture. One-way ANOVA was used and bars marked with an asterisk were significantly higher than LB (Dunnett’s test, p < 0.05).

Growth of PA2 in cell-free supernatants of 0.1229 also amplified Stx2a production, indicating this phenomenon does not require whole cells (Fig. 2A). Sequencing of the genome of 0.1229 using Illumina technology revealed that it belonged to the same sequence type (ST73) as *E. coli* strains CFT073 (27) and Nissle 1917 (28), and carried a plasmid similar to pRS218 in *E. coli* RS218 (29). However, supernatants harvested after growth of these strains failed to increase Stx2a production by PA2 (Fig. 2A). To test whether increased Stx2a production was dependent on *recA*, 0.1229 was co-cultured with the W3110Δ*tolC* P*recA-gfp* reporter strain. As anticipated, we found that among this collection, only strain 0.1229 increased GFP expression as a co-culture (Fig. 2B). Treatment of 0.1229 supernatants revealed this bioactivity was resistant to boiling, and sensitive to Proteinase K (Fig. 2C).

**Fig. 2:**
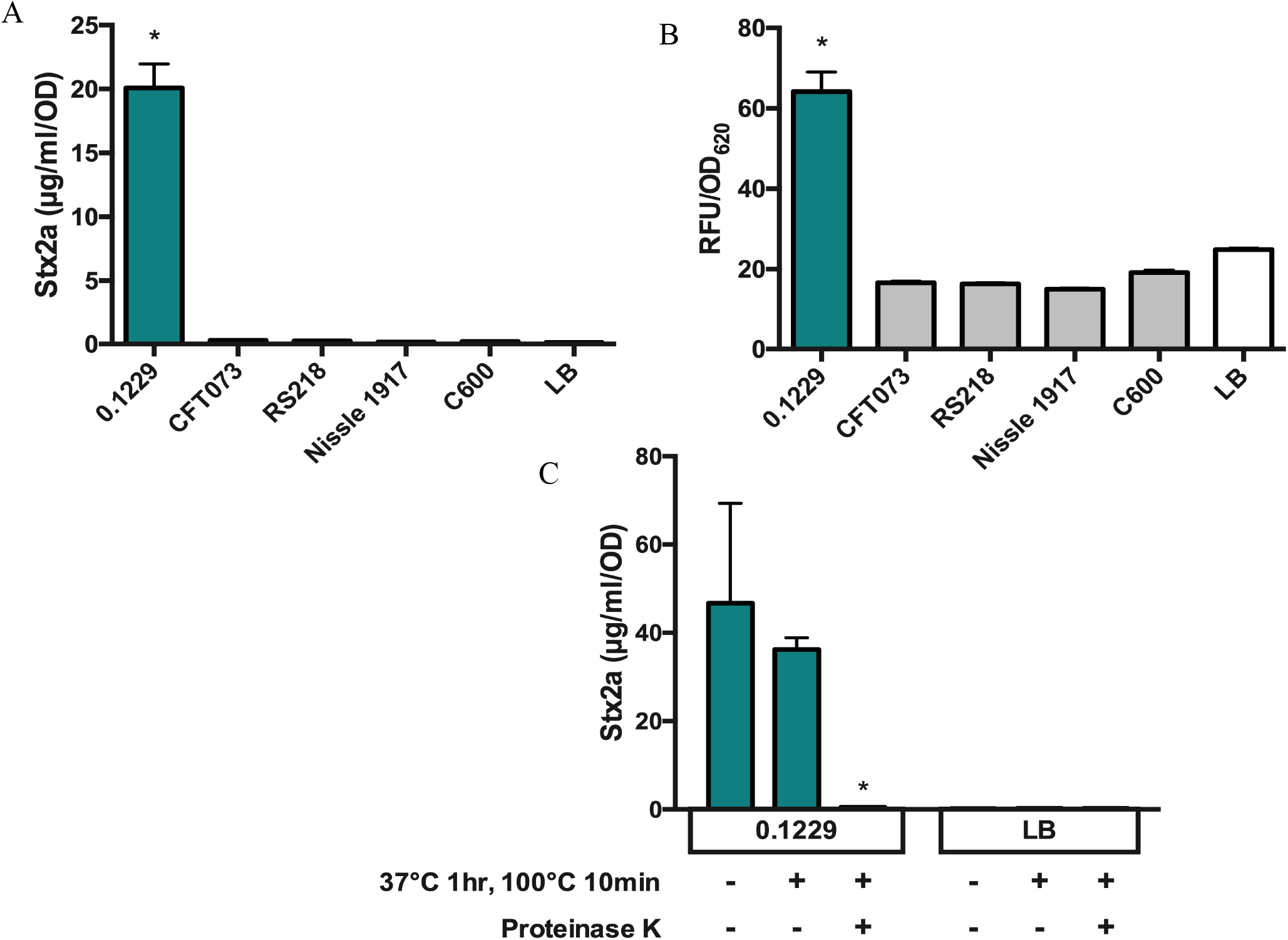
The Stx2a levels (A) and fluorescence (B) of non-pathogenic *E. coli*, after PA2 growth in cell-free supernatant or W3110Δ*tolC* P*recA-gfp* co-culture, respectively. Samples were normalized to cell density, OD_600_ or OD_620_, for Stx2a or fluorescence, respectively. One-way ANOVA was used and levels marked with an asterisk were significantly higher than LB (Dunnett’s test, p < 0.05). The Stx2a levels (C) of PA2 grown in 0.1229 cell-free supernatant (2C-left) or LB (2C-right) with or without heat and Proteinase K treatments. Two-way ANOVA was used and bars marked with an asterisk were significantly lower than untreated 0.1229 or LB (Dunnett’s test, p < 0.05).

### The plasmids of 0.1229 may play a role in Stx2a amplification

Further analysis of the Illumina sequence data revealed high sequence identity between the chromosomes of 0.1229, CFT073, Nissle 1917 and RS218 (data not shown). The most notable differences were in predicted plasmid content. To obtain a more complete picture, PacBio long read technology was used to sequence the genome of 0.1229.

The largest plasmid of 0.1229, designated p0.1229_1, was 114,229 bp, and 99.99% identical with 100% query coverage to pUTI89 (30) and pRS218 (29) (Fig. 3A). A second plasmid designated p0.1229_2 had five identifiable antimicrobial resistance genes, was 96,272 bp, and encoded the operon for microcin B17 (MccB17), a microcin that inhibits DNA gyrase (31). The plasmid p0.1229_2 shared high sequence identity with other known plasmids, being 99.96% identical with 57% query to pRS218, 97.81% identical with 82% query to pECO-fce (NCBI accession CP015160) and 100% identical with 89% query to pSF-173-1 (32) (Fig. 3B). A third plasmid was smaller than the cutoff for size selection used during PacBio library preparation but was closed by Illumina sequencing. This plasmid, designated p0.1229_3, was 12,894 bp and encoded ampicillin resistance (*blaTEM-1b*) (Fig. 3C). It was similar to pEC16II (NCBI accession KU932034), 99.85% identical with 61% query, and to pHUSEC41-3 (33), 98.89% identical with 61% query.

**Fig. 3:**
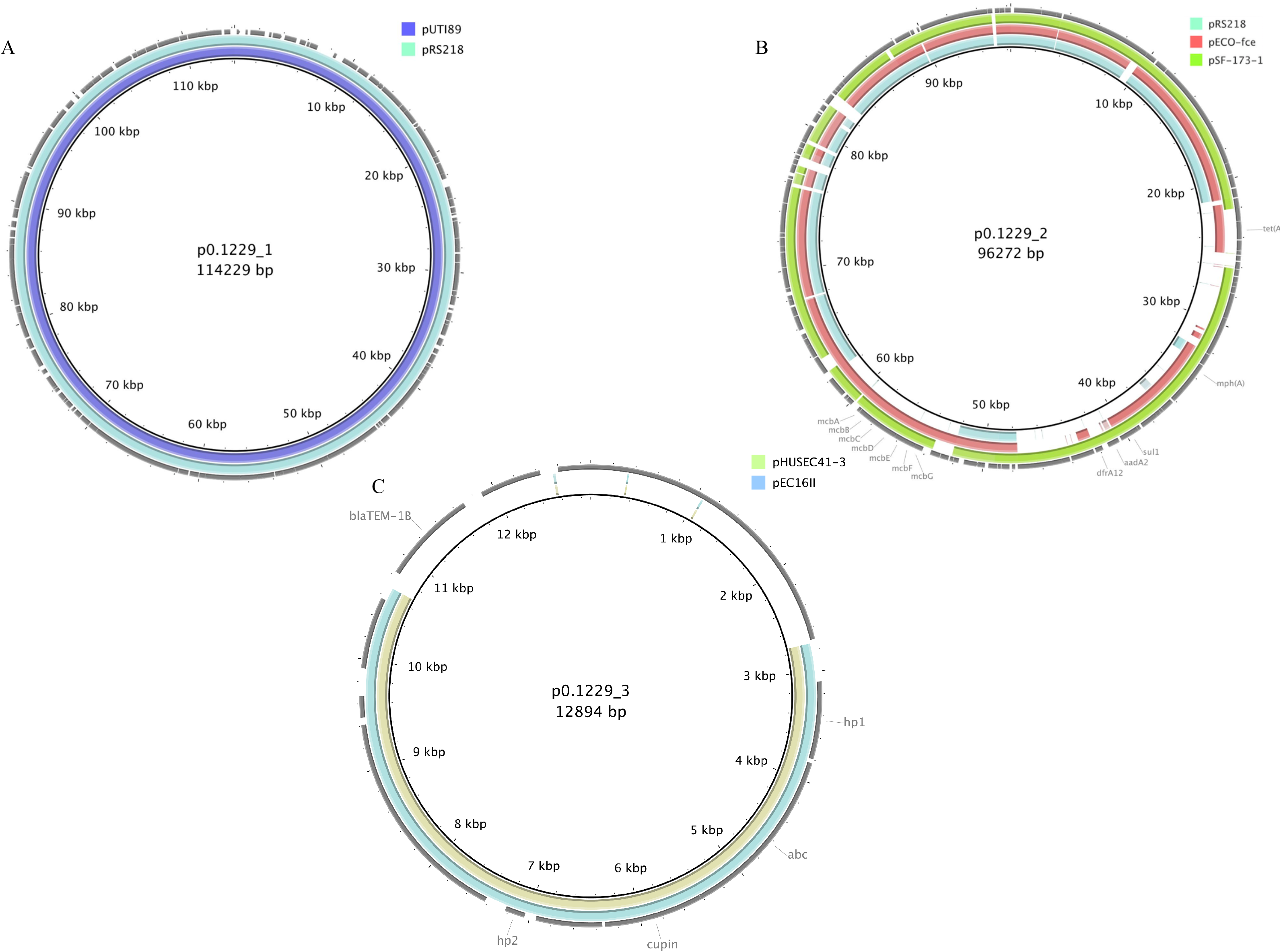
Plasmid comparisons of individual 0.1229 plasmids and publicly available plasmids from NCBI by BLAST and visualized with BRIG. The colored rings indicate sequence similarity and the outer grey ring denotes annotated ORFs. The MccB17 operon is labeled *mcbA-mcbG*, and five antimicrobial resistance genes, macrolide (*mph(A)*), tetracycline (*tet(A)*), sulphonamide (*sul1*), aminoglycoside (*aadA2*), and trimethoprim (*dfrA12*) were identified (B). Ampicillin resistance gene (*blaTEM-1B*) and four ORFs, *hp1, abc, cupin* and *hp2* are also labeled (C). NCBI accession numbers are pUTI89 (CP000244), pRS218 (CP007150), pECO-fce (CP015160), pSF-173-1 (CP012632), pHUSEC41-3 (NC_018997), and pEC16II (KU932034).

As RS218 did not amplify Stx2a production (Fig. 2A), it was assumed that p0.1229_1 did not encode the genes responsible for this phenotype. Similarly, *E. coli* strain SF-173 did not increase GFP when co-cultured with W3110Δ*tolC* P*recA-gfp*, suggesting genes encoded on p0.1229_2 were also not required (data not shown).

### Strain 0.1229 encodes microcin B17, which is partially responsible for Stx2a amplification

Microcin B17 (MccB17) is a 3.1 kDa (43 amino acid) DNA gyrase inhibitor that is found on a seven gene operon, with *mcbA* encoding the 69 amino acid microcin precursor (31). Although pSF-173-1 encodes this operon, there was a three-nucleotide deletion observed in *mcbA* in pSF-173-1, compared to an earlier published sequence (31). This deletion is predicted to shorten a ten Gly homopolymeric stretch by one amino acid residue. Although this Gly rich region is not important for interaction with the gyrase-DNA complex (34), it seemed prudent to confirm that the results reported above with strain SF-173 were not due to production of a non-functional McbA. Therefore, knockouts of *mcbA* (Δ*mcbA*), and the entire operon (Δ*mcbABCDEFG*) were constructed in 0.1229. These mutations decreased Stx2a amplification by O157:H7 compared to wildtype 0.1229 (Fig. 4A) however they did not ablate the Stx2a levels back to mono-culture levels. Similar results were seen with the P*recA-gfp* strain (Fig. 4B), although differences were less pronounced than those seen with the Stx assays.

**Fig. 4:**
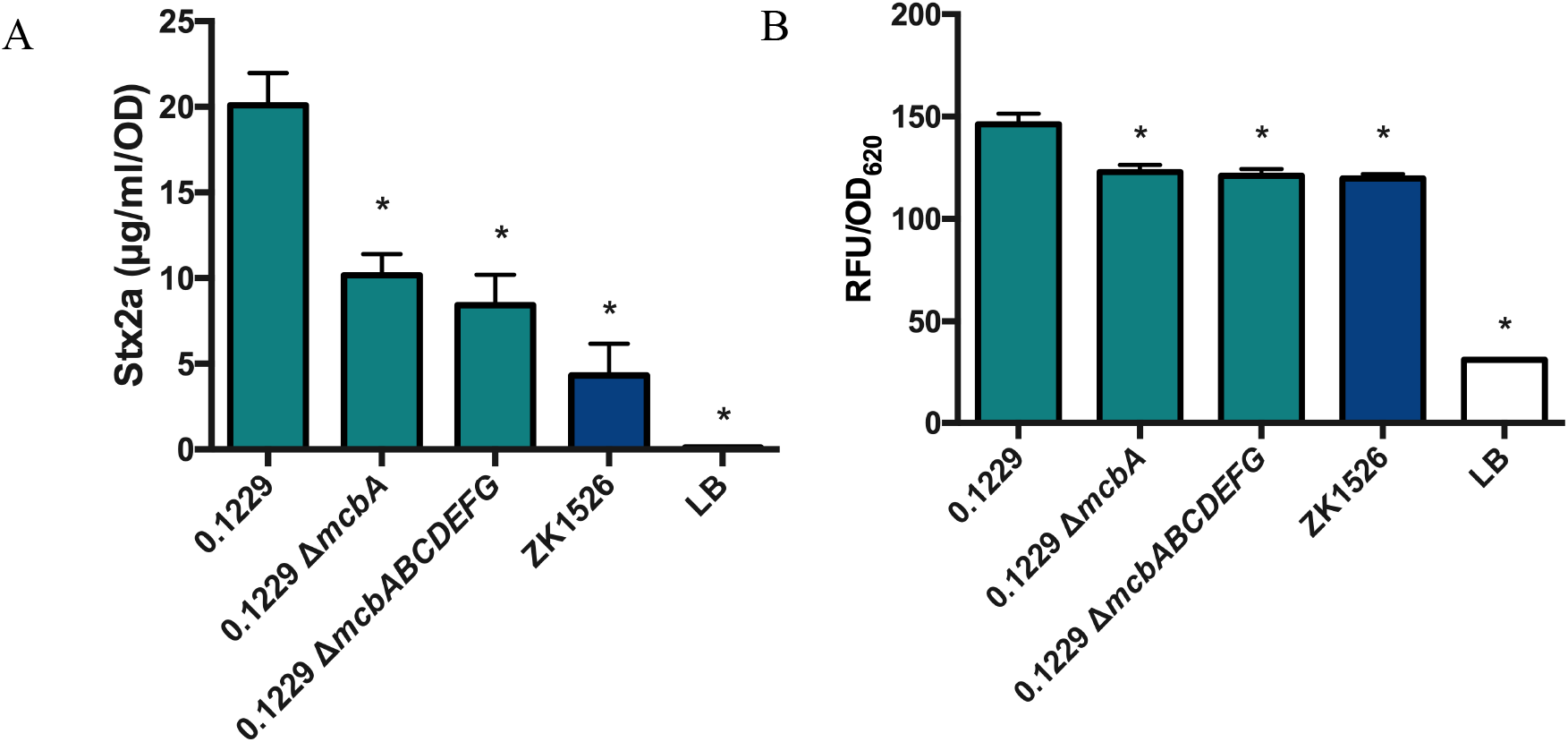
The Stx2a levels (A) and fluorescence (B) of 0.1229, its MccB17 knockouts and ZK1526, after PA2 growth in cell-free supernatant or W3110Δ*tolC* P*recA-gfp* co-culture, respectively. Samples were normalized to cell density, OD_600_ or OD_620_, for Stx2a or fluorescence, respectively. One-way ANOVA was used and bars marked with an asterisk were significantly lower than 0.1229 (Dunnett’s test, p < 0.05).

### Four ORFs encoded by p0.1229_3 are necessary for Stx2a amplification phenotype

It was next hypothesized that p0.1229_3 encoded the activity responsible for Stx2a amplification by 0.1229Δ*mcbA* and 0.1229Δ*mcbABCDEFG*. A C600 strain transformed with p0.1229_3 amplified Stx2a production of PA2 (Fig. 5), confirming the importance of this plasmid. By systematically deleting portions of p0.1229_3, two regions were identified as essential for increased Stx2a production (Fig. 6A). The genes annotated in these regions are referred to as hypothetic proteins (hp), domains of unknown (DUF), an ATP-binding cassette (ABC)-type transporter and a member of the cupin superfamily of conserved barrel domains. The mutant, 0.1229 Δ6 (2850-5473 bp), deleted two open reading frames (ORFs), referred to as *hp1* and ABC, and 0.1229 Δ7 (5426-7950 bp) deleted *cupin, DUF4440, DUF2164, hp2, hp3* and a portion of a nuclease (Fig. 6B). These results were confirmed using co-culture assays with the P*recA-gfp* reporter (Fig. S1). Insertional inactivation of individual ORFs in these regions identified four, *hp1, abc, cupin*, and *hp2*, that were necessary for enhanced Stx2a production (Fig. 6C). Similar results were shown in co-culture with P*recA-gfp*, although 0.1229Δ*hp2* showed only a moderate decrease in GFP expression (Fig. S1). Cloning of a 5.2kb region of the plasmid, spanning upstream of *hp1* through the beginning of the putative nuclease-encoding gene, confirmed this activity is encoded within this region (Fig. 6D). Cloning of a similar region that ended after *abc* did not provide C600 the ability to increase GFP (data not shown).

**Fig. 5:**
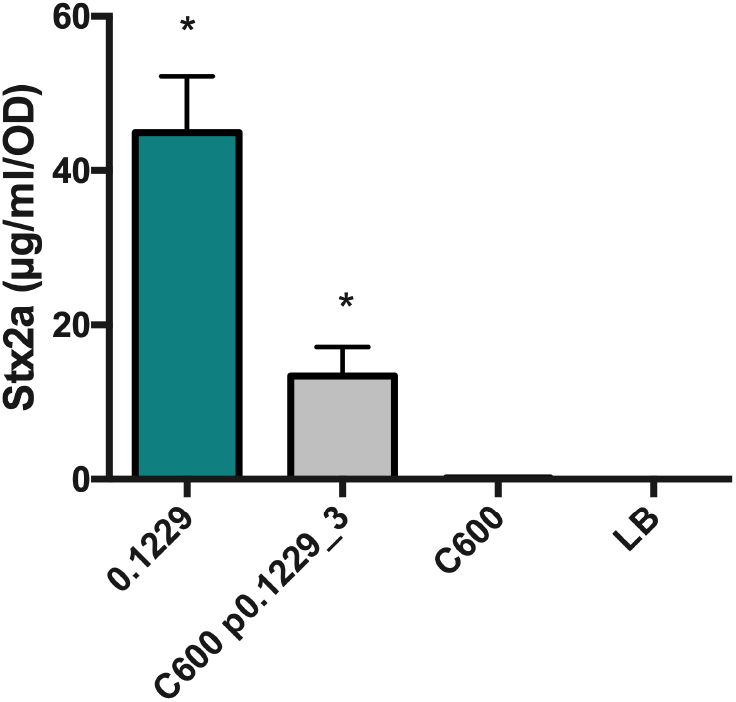
PA2 was grown in the cell-free supernatant of 0.1229, C600 containing p0.1229_3 and C600. Stx2a levels were measured using the R-ELISA. LB refers to PA2 grown in LB broth. One-way ANOVA was used and bars marked with an asterisk were significantly higher than LB (Fisher’s LSD test, p < 0.05).

**Fig. 6:**
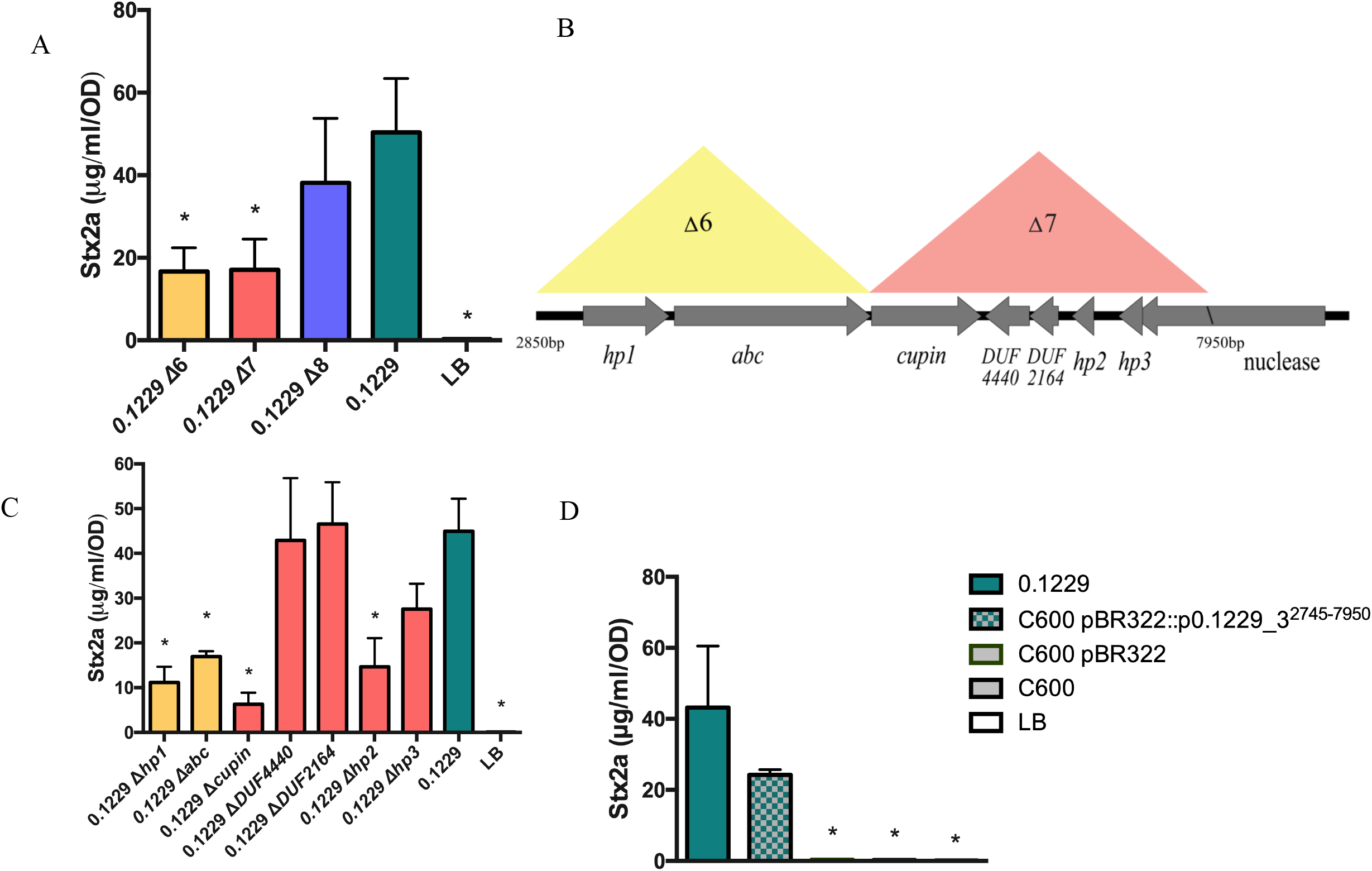
PA2 was grown in the cell-free supernatant of 0.1229 knockouts (A & C). Portion of p0.1229_3 is depicted with predicted open reading frames (ORFs) (B). The colored triangles indicate the name of the regional knockout., PA2 grown in the supernatant of a C600 strain containing a portion of p0.1229_3 (pBR322::p0.1229_3^2745-7950^) (C). Stx2a levels measured using the R-ELISA. LB refers to PA2 grown in LB broth. One-way ANOVA was used and bars marked with an asterisk were significantly lower than 0.1229 (Fisher’s LSD test, p < 0.05).

*In silico* comparisons identified a nearly identical gene cluster in other species, including *Shigella sonnei* and *Klebsiella pneuomoniae* (Fig. S2). The region of p0.1229_3 spanning nucleotides 2745 to 7238bp was greater than 99.6% identical on the nucleotide level, when comparing all the strains in Fig. S2. Similar gene clusters containing Hp1 at 36 to 68% amino acid identity, were found in other species as well (Fig. S3). In these clusters, orthologs to *hp1, abc*, and *cupin* were commonly co-localized and the genes were found in the same order.

### The secreted molecule requires *tolC* for secretion, and *tonB* for import into target strains

Some bacteriocins and microcins require genes encoded outside of the main operon for secretion, such as the efflux protein TolC (35). The supernatant of 0.1229Δ*tolC* did not increase Stx2a expression by strain PA2 to levels seen with wildtype 0.1229 supernatants (data not shown). Similar results were observed in co-culture experiments using the P*recA-gfp* carrying strain (Fig. 8). The phenotype was restored when *tolC* was complemented on a plasmid, but only when tested with the P*recA-gfp* strain (Fig. 7). Similarly, numerous bacteriocins are translocated into target cells using the TonB system (36). A *tonB* knockout was constructed in the P*recA-gfp* reporter strain, as we were unsuccessful generating this in a O157:H7 background. In co-culture with 0.1229, the MG1655Δ*tonB* P*recA-gfp* strain produced lower GFP levels than the MG1655 P*recA-gfp* strain (Fig. 8). This phenotype was restored when pBAD24::*tonB*, but not pBAD24, was transformed into the mutant strain (Fig. 8).

**Fig. 7:**
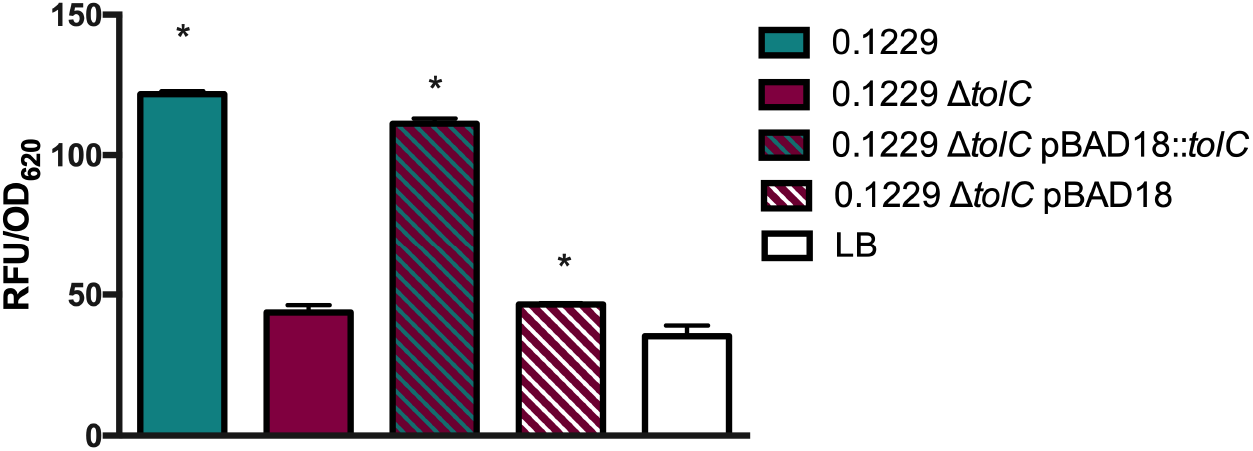
W3110Δ*tolC* P*recA-gfp* was grown in co-culture with 0.1229, 0.1229Δ*tolC* and 0.1229Δ*tolC* pBAD18 containing strains, and fluorescence was measured. Samples were normalized to cell density, OD_620_. One-way ANOVA was used and bars marked with an asterisk were significantly higher than LB (Dunnett’s test, p < 0.05).

**Fig. 8:**
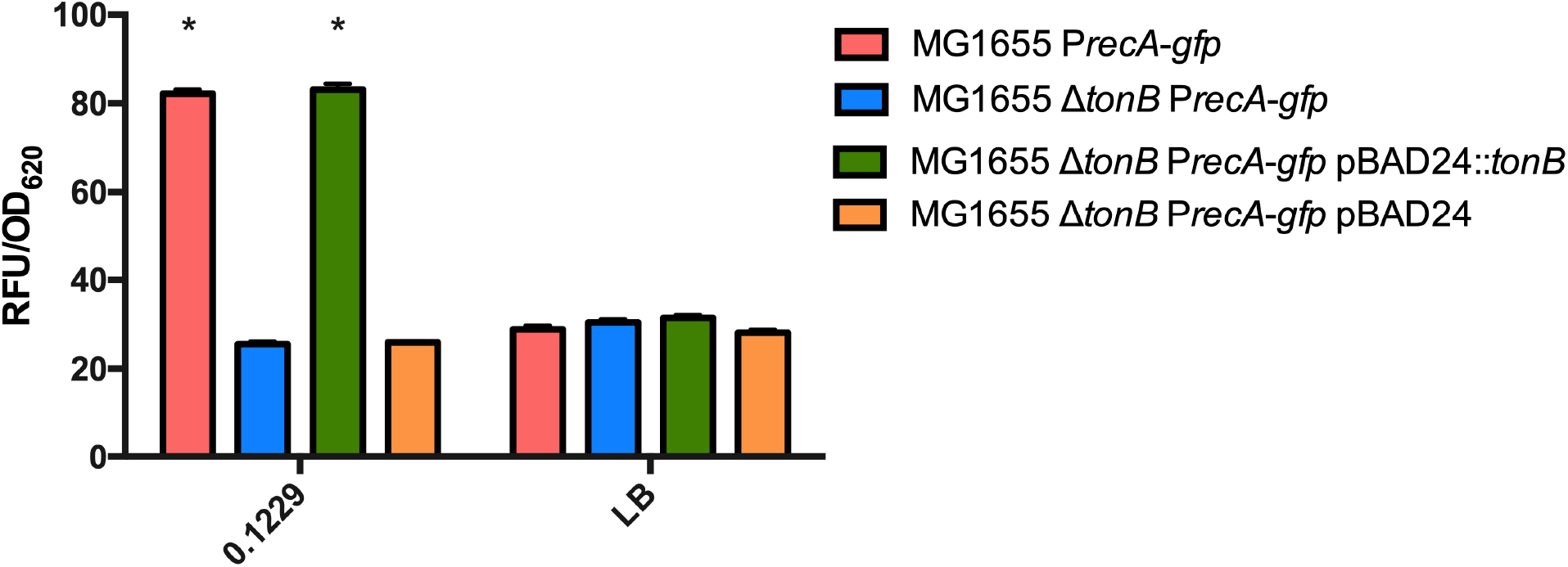
Plasmid expressing P*recA-gfp* was electroporated into MG1655, MG1655Δ*tonB*, MG1655Δ*tonB* pBAD24 and MG1655Δ*tonB* pBAD24::*tonB*. These strains were grown in coculture with 0.1229, or by themselves (LB). Two-way ANOVA was used and bars marked with an asterisk were significantly higher than their respective monoculture control (Dunnett’s test, p < 0.05).

### The gene cluster was identified in additional strains

Lastly, it was hypothesized that *E. coli* isolated from human feces would encode the similar molecules identified here. A total of 101 human fecal *E. coli* isolates were obtained from Penn State’s *E. coli* Reference Center, and three of these were found to induce GFP production in the P*recA-gfp* reporter assay (Fig. S4). Furthermore, the supernatants of two of these isolates, designated 91.0593 and 99.0750, increased Stx2a to levels similar to 0.1229, however 90.2723 did not (Fig. 9A). Genome sequencing of these three organisms revealed that 91.0593 and 99.0750 carried plasmids similar to p0.1229_3, however the latter plasmid had a deletion in the recombinase and transposon regions (Fig. 9B). Strain 99.0750 was molecular serotype O36:H39, while 91.0593 could not be O typed but was identified as H10.

**Fig. 9:**
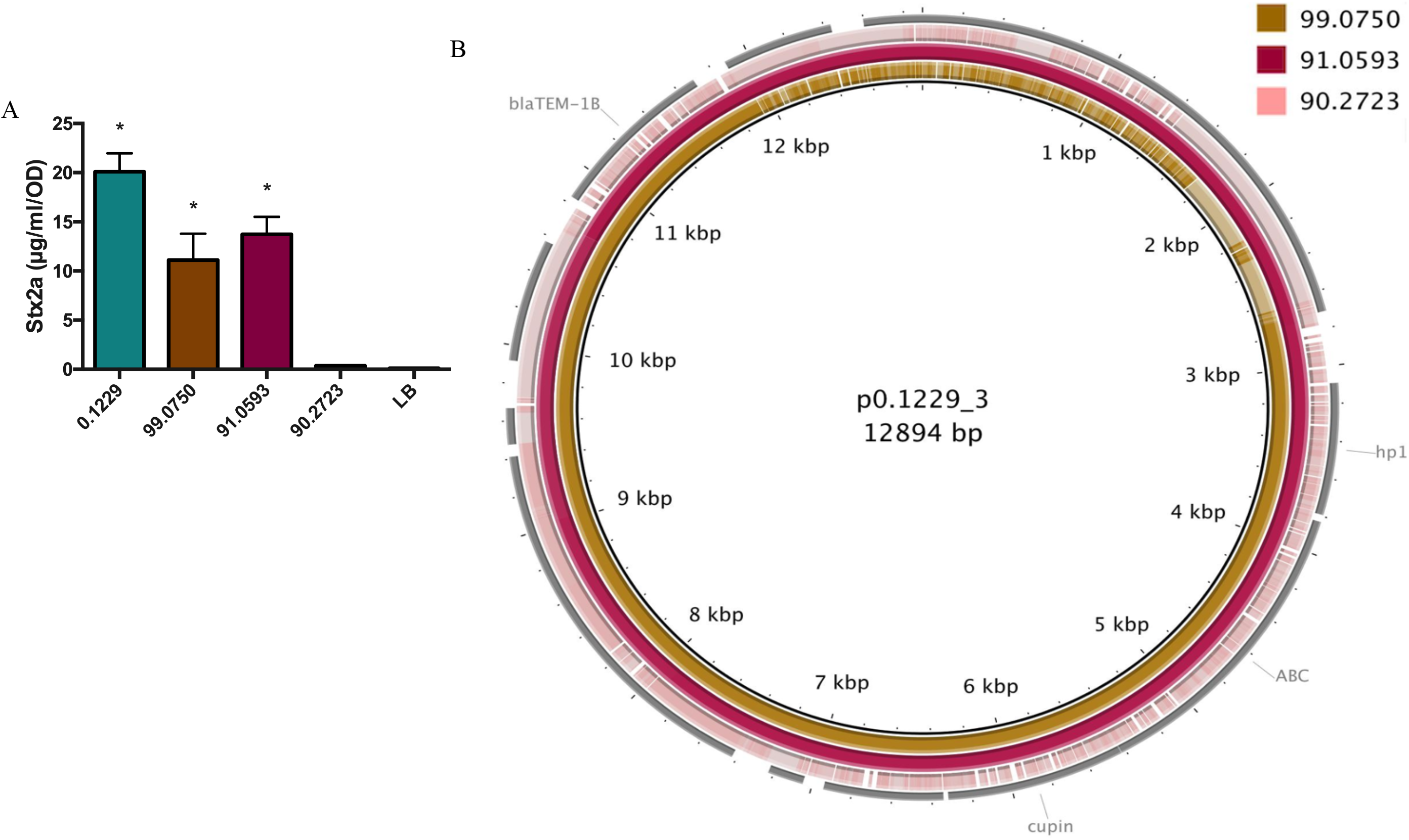
PA2 was grown in the cell-free supernatant of 0.1229 and three human fecal *E. coli* isolates (A). Stx2a levels measured using the R-ELISA. LB refers to PA2 grown in LB broth. One-way ANOVA was used and bars marked with an asterisk were significantly higher than LB (Dunnett’s test, p < 0.05). p0.1229_3 was compared to the contigs of 99.0750, 91.0593 and 90.2723 using BLAST and visualized using BRIG (B).

## Discussion

The concentration of *E. coli* in human feces ranges from 10^7^ to 10^9^ colony forming units (CFU) (37, 38). Typically, there are up to five commensal *E. coli* strains colonizing the human gut at a given time (39, 40). As the human microbiota affects O157:H7 colonization and virulence gene expression (41–44), it is thought that community differences in the gut microflora may explain, in part, individual differences in disease symptoms (45). Indeed, commensal *E. coli* that are susceptible to *stx2*-converting phage can increase phage and Stx production (22, 23). In mice given a co-culture of O157:H7 and phage-resistant *E. coli*, minimal toxin was recovered in the feces, but with *E. coli* that were phage-susceptible, higher levels of toxin were found (46). However, it is clear that phage infection of susceptible bacteria is not the only mechanism by which the gut microflora affects Stx2 levels during infection (19, 20, 24, 26).

In this study, both whole cells and spent supernatants of *E. coli* 0.1229 enhanced Stx2a production by *E. coli* O157:H7 strain PA2. This latter strain is a member of the hypervirulent clade 8 (47) and was previously found to be a high Stx2a producer in co-culture with *E. coli* C600 (23). *E. coli* 0.1229 produces at least two molecules capable of increasing Stx2a. The first is MccB17, a DNA gyrase inhibitor, shown to activate Stx2a production in an earlier study (24). This current study identified a second molecule localized to a 12.8 kb plasmid, and all genes necessary for production are found within a 5.2 kb region. Furthermore, gene knockouts identified four potential ORFs within this region, *hp1, abc, cupin* and *hp2*, that are required for 0.1229 mediated Stx2a amplification. This gene cluster was also identified on pB51 (48), a similar plasmid to p0.1229_3, however limited characterization was reported.

Oxidizing agents, such as hydrogen peroxide (H_2_O_2_), and antibiotics targeting DNA replication, such as ciprofloxacin, mitomycin C and norfloxacin, are known to induce *stx*-converting phage (5, 49, 50) and subsequently Stx2 production (5, 49). However, the Stx2 amplifying activity of the 0.1229 supernatant was abolished by Proteinase K, suggesting the inducing molecule is proteinaceous in nature. Colicins are bacteriocins found in *E. coli* (51), are generally greater than 30 kDa in size, and at least one member has been previously shown to enhance O157:H7 Stx2 production (26). While some colicins utilize TonB for translocation, they are not expected to be heat stable. The molecule produced by 0.1229 was resistant to 100°C for 10 minutes, strongly suggesting it is not a colicin.

Microcins are bacteriocins that are generally smaller than 10 kDa. Their size and lack of secondary and tertiary structure make them more heat stable than colicins. Microcins are divided into three classes; class I and class IIa are plasmid encoded, while class IIb are chromosomally encoded. Class I and IIb are post-translationally modified (52, 53), while class IIa are not. To date, all class II but only one member of class I (microcin J25) use an ATP-binding cassette (ABC)-type transporter in complex with TolC for export (35), and the TonB system for import into target cells (36). The putative microcin produced by 0.1229 is plasmid encoded, along with a predicted ABC transporter and is TolC and TonB dependent. Therefore, this microcin appears to be more closely related to class IIa microcins. However, purification of the microcin to identify possible post translational modifications is necessary to confirm whether designating as class I or IIa is more appropriate.

There are four known class IIa microcins, microcin V (MccV, previously named colicin V) (54, 55), microcin N (MccN, previously named Mcc24) (56), microcin L (MccL) (57), and microcin PDI (MccPDI) (58, 59). The operons encoding these microcins contain four or five genes, including the microcin precursor, immunity and export genes. MccN also encodes a putative regulator, with a histone-like nucleoid domain (56). The microcin precursor genes possess leader sequences of approximately 15 amino acids, containing the signature sequence MRXI/LX(9)GG/A (X=any amino acid), and are typically cleaved by the ABC transporters during export (60). A potential leader sequence with the double glycine was found in *hp2*. Additionally, a small peptide (DHGSR) was identified in the supernatants of 0.1229 by mass spectroscopy (data not shown) corresponding to an ORF internal to *hp2* encoded in the opposite direction. Future experiments will determine if one of these, or another region, encodes a secreted microcin.

One argument against designation as a class IIa microcin, is the lack of an identifiable N-terminal proteolytic domain (61) in the predicted ABC transporter encoded on p0.1229_3. This domain is found in all other members of class IIa. Interestingly, the class I microcin J25 (MccJ25) also encodes an ABC transporter lacking this domain. Unlike the other class I microcins, MccJ25 is TolC and TonB dependent for export and import, respectively. While the possibility cannot be excluded that the system identified here is a class I microcin, if so, it is more similar to MccJ25 than to other members of this group.

While the current mechanism of action is unknown, it is theorized that the putative microcin causes DNA damage, through double strand breaks, depurination, or inhibition of DNA replication. Such actions would lead to RecA-dependent phage induction and Stx2 production. The suspected mode of action would be divergent from the known class IIa microcins, which target the inner membrane (62) and MccJ25 which inhibits the RNA polymerase (63). Besides the predicted ABC transporter, the functions of the other ORFs is unclear. We anticipate one of these may encode an ABC accessory protein, known to be essential for these export complexes (64). One ORF encodes a cupin domain found in a functionally diverse set of proteins. An immunity gene protecting the host may also be expected in this region.

The genes encoding the putative microcin were additionally found in *E. coli* strains 99.0750 and 91.0593. Genome sequencing of these strains failed to identify genes encoding MccB17, which may explain the lower levels of Stx2a production seen in co-culture with PA2 compared to those seen with 0.1229. Bioinformatic analyses also identified other *E. coli* that encode nearly identical regions. Interestingly, one of these was *E. coli* O104:H4 HUS, isolated in 2001 (33), and responsible for a large 2011 outbreak in Germany. However, a premature stop codon identified in *cupin* suggests it is non-functional. Homologs of *hp1, ABC* and *cupin* were identified together in several other organisms distantly related to *E. coli*, suggesting these encode a functional unit. The absence of *hp2* in most of these genetic clusters argues against this ORF encoding the anti-bacterial activity or may suggest that these organisms encode microcins distinct from *hp2*.

In conclusion, a putative microcin was identified in *E. coli*, expanding our knowledge of this small group of antimicrobial peptides. This study also identifies another mechanism by which *E. coli* may enhance Stx2a production by *E. coli* O157:H7. Further studies may also provide new insights into the diverse genetic structure and functions of microcin-encoding systems.

## Materials & Methods

### Bacterial strains, media and growth conditions

*E. coli* strains were grown in Lysogeny Broth (LB) at 37°C unless otherwise indicated, and culture stocks were maintained in 20% glycerol at −80°C. Antibiotics were used at the following concentrations; ampicillin (100 μg/ml), chloramphenicol (25 μg/ml), kanamycin (50 μg/ml), and tetracycline (10 μg/ml). All bacterial isolates, plasmids and primers used in this study can be found in Table 1. *E. coli* SF-173-1 was provided by Dr. Craig Stephens, Santa Clara University.

**Table 1:**
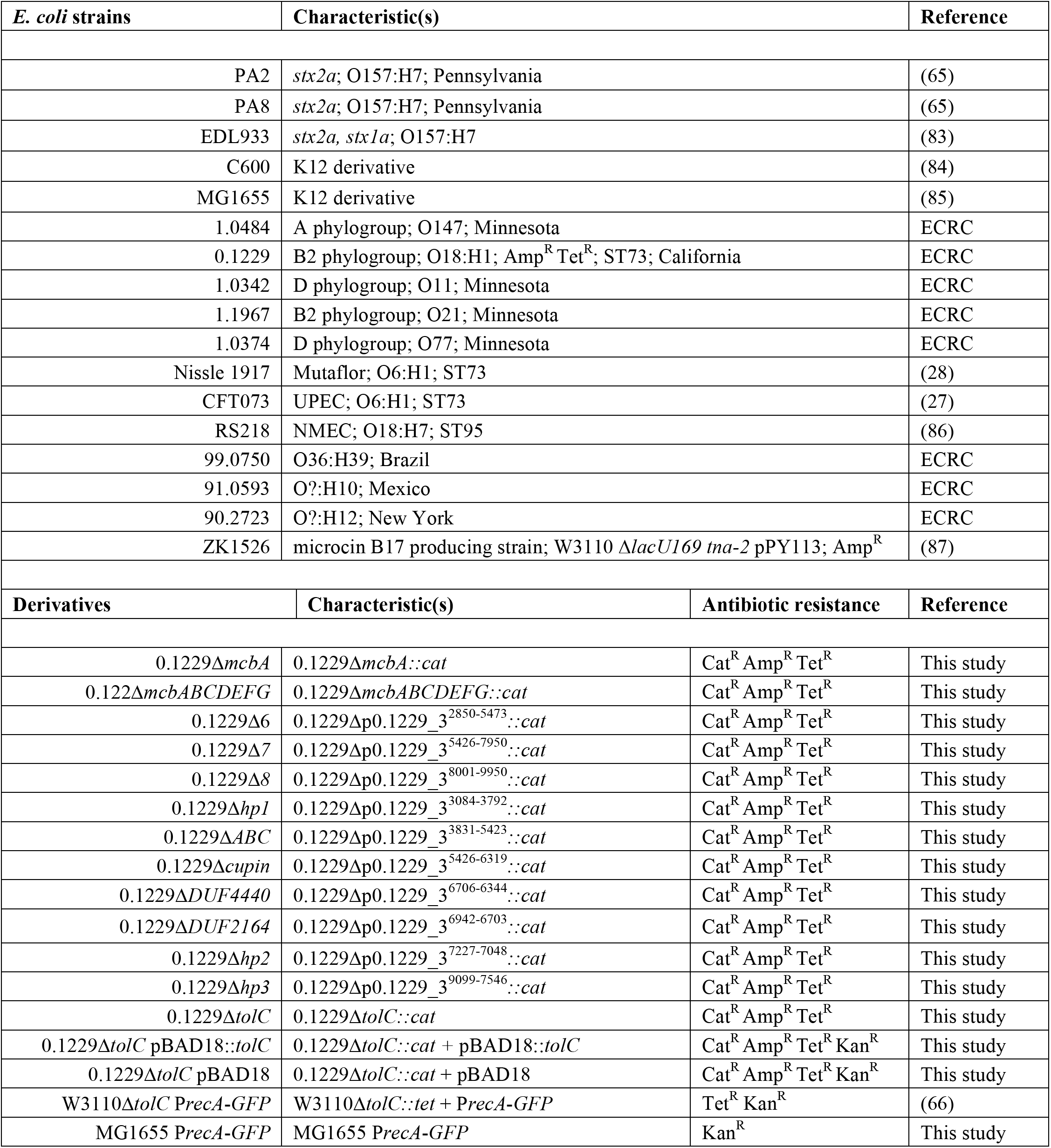
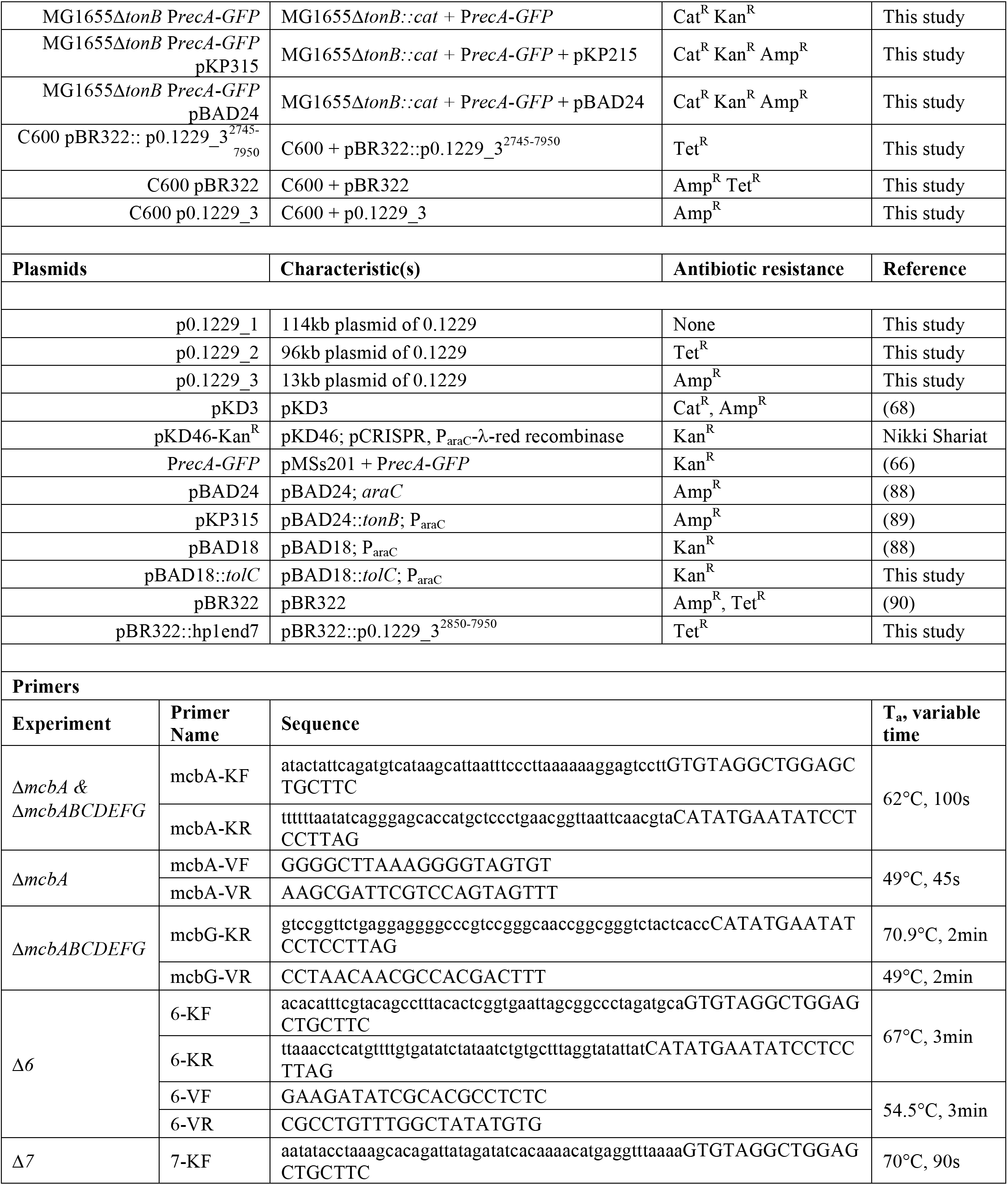
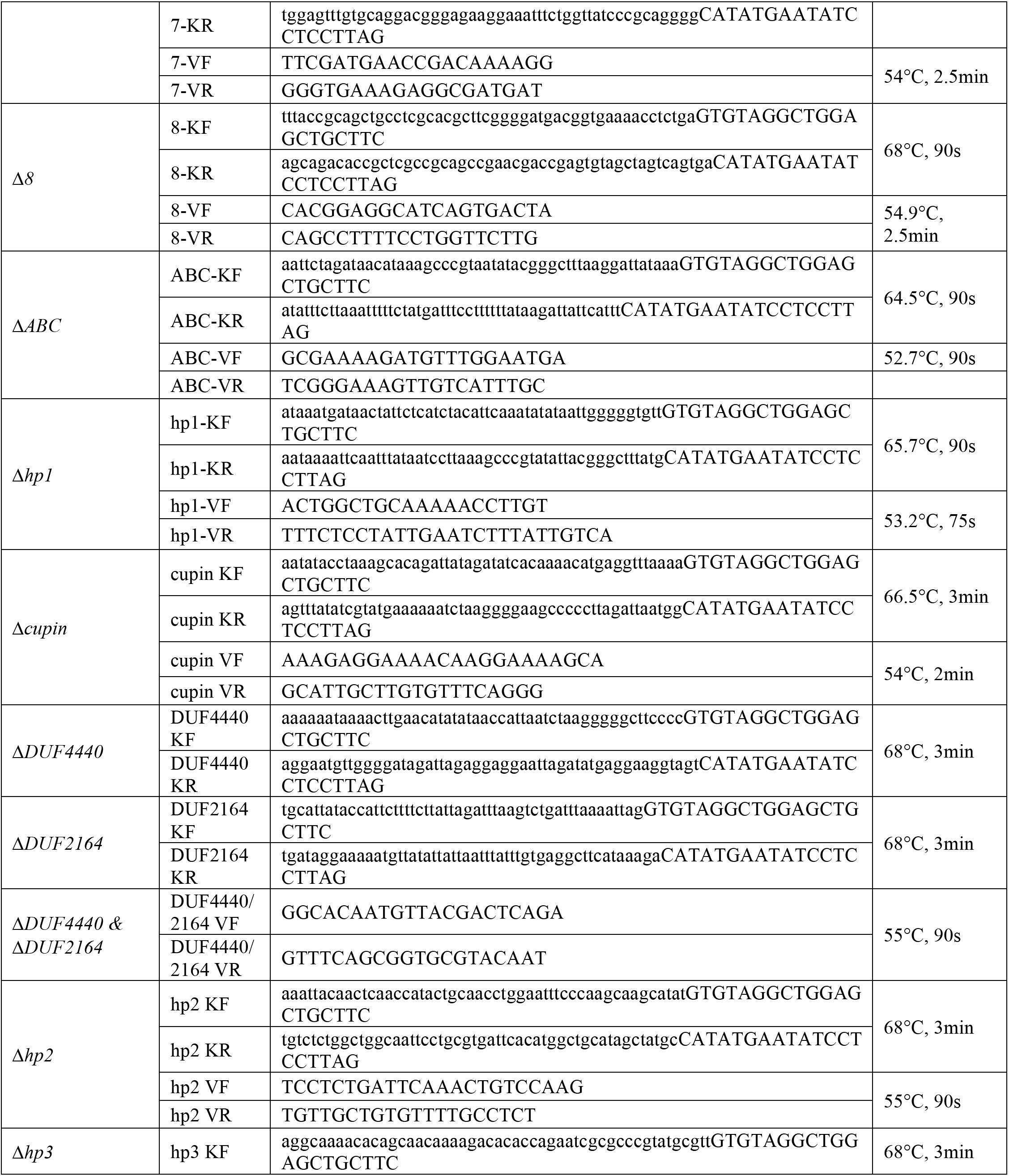
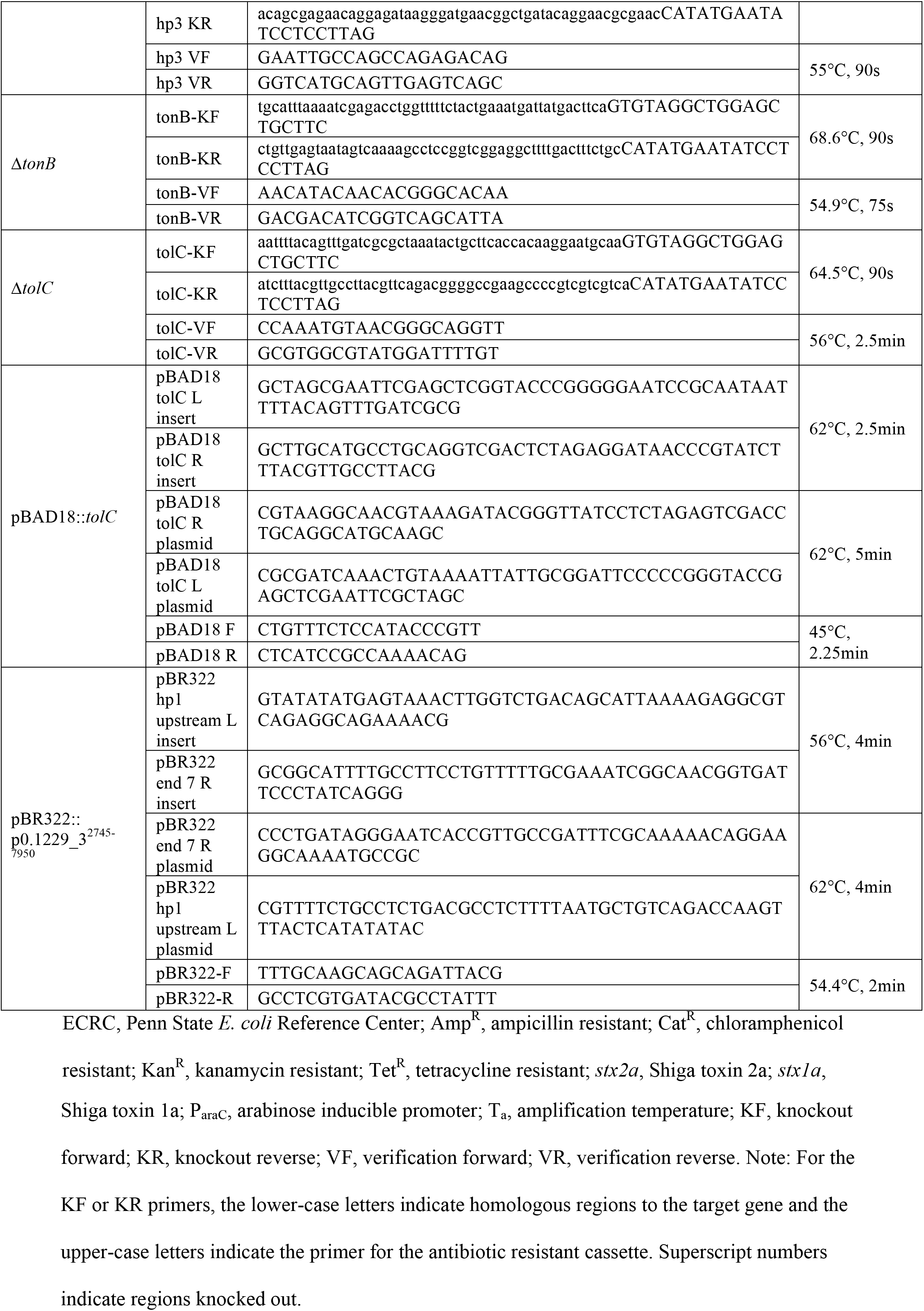
Bacterial isolates, plasmids and primers used in this study

### Co-culture with PA2

Co-culture with *E. coli* O157:H7 PA2 was performed similar to previously described (23). PA2 and commensal *E. coli* strains were grown overnight at 37°C (with shaking at 250 rpm). LB agar (2.5 ml) was added to 6-well plates (BD Biosciences Inc., Franklin Lakes, NJ), and allowed to solidify. PA2 and commensal strains were each diluted to an OD_600_ of 0.05 in 1 ml of LB broth and added to the 6-well plates. A mono-culture of PA2 (at 0.05 OD_600_ in 1ml) served as a negative control. The plates were incubated without shaking at 37°C. After 16 hr, cultures were collected, cells were lysed with 6 mg/ml polymyxin B at 37°C for 5 min, and supernatants were collected. Samples were immediately tested with the receptor-based enzyme-linked immunosorbent assay (R-ELISA), as described below, or stored at −80°C. Total protein was calculated using the Bradford assay (VMR Life Science, Philadelphia, PA), and used to calculate μg/mg Stx2.

### R-ELISA for Stx2a detection

Detection of Shiga toxin was performed using a sandwich ELISA approach, previously described by Xiaoli *et al*., 2018 (24). Briefly, 25 μg/ml of ceramide trihexosides (bottom spot) (Matreya Biosciences, Pleasant Gap, PA) dissolved in methanol was used for coating of the plate. Washes were performed between each step using PBS and 0.05% Tween-20. Stx2a-containing samples were diluted in PBS as necessary to obtain final readings in the linear range. Samples were added to the wells in duplicate and incubated with shaking for 1 hr at room temperature. Supernatants of *E. coli* PA11, a high Stx2a producer (65), were used as a positive control. Anti-Stx2 monoclonal mouse antibody (Santa Cruz Biotech, Santa Cruz CA) was added to the plate at a concentration of 1 μg/ml, then incubated for 1 hr. Anti-mouse secondary antibody (MilliporeSigma, Burlington MA) conjugated to horseradish peroxidase (1 μg/ml) was added to the plate, and incubated for 1 hr. For detection, 1 step Ultra-TMB (Thermo-Fischer, Waltham, MA) was used, and 2M H_2_SO_4_ was added to the wells to stop the reaction. The plate was read at 450 nm using a DU®730 spectrophotometer (Beckman Coulter, Atlanta, GA). A standard curve was generated from two-fold serially diluted PA11 samples and used to quantify the μg/ml of Stx2a present in each sample.

### Cell-free supernatant assay with PA2

*E. coli* O157:H7 strain PA2 and non-pathogenic *E. coli* strains were individually grown with shaking at 37°C for 16 hr. Overnight culture of the non-pathogenic strains were centrifuged, and supernatants were filtered through 0.2 μm cellulose filters (VWR International, Radnor, PA). LB agar (2.5 ml) was added to the wells of 6-well plates (BD Biosciences Inc., Franklin Lakes, NJ) and allowed to solidify. PA2 was added to wells at a final density of 0.05 OD_600_ in 1 ml of spent supernatant. For the negative control, PA2 was resuspended in fresh LB broth to the same cell density, and 1 ml was added to a well. The plates were statically incubated at 37°C for 8 hr, after which the cell density (OD_600_) was recorded. Cells were lysed with 6 mg/ml Polymyxin B at 37°C for 5 min and supernatant recovered. Samples were immediately tested for Stx2a by R-ELISA or stored at −80°C. Data reported as μg/ml/OD_600_.

### Detection of SOS inducing agents using P*recA-gfp*

*E. coli* expressing P*recA-gfp*, which encodes green fluorescent protein (*gfp*) under control of the *recA* promoter (66), was purchased from Dharmacon (Lafayette, CO). The plasmid was transformed into *E. coli* W3110Δ*tolC*. The *tolC* deletion reduces the potential efflux of *recA*-activating molecules. W3110Δ*tolC* P*recA-gfp* and commensal strains were individually grown overnight with shaking at 37°C. LB agar (2.5 ml) was added to 6-well plates and allowed to solidify. W3110Δ*tolC* P*recA-gfp* and one commensal strain were each diluted to a final OD_600_ of 0.05 in LB broth, and 1ml was added to the 6-well plates. The negative control included only W3110Δ*tolC* P*recA-gfp* at a final OD of 0.05 in 1 ml LB broth. The plates were statically incubated at 37°C. After 16 hr, 100 μl was removed from each well, added to black 96 well clear bottom plates (Dot Scientific Inc., Burton, MI) and optical density (OD_620_) was read using a DU®730 spectrophotometer. Relative fluorescence units (RFU) were measured at an excitation of 485 nm and emission of 538 nm on a Fluoroskan Ascent FL (Thermo Fisher Scientific, Waltham, MA) (67). RFU values were normalized to cell density.

### One step recombination for *E. coli* knockouts

Mutants of 0.1229 and MG1655 were constructed using one-step recombination (68). Primers contained either 50 bp upstream or downstream of the gene of interest, followed by sequences annealing to the P1 and P2 priming sites from pKD3. PCR was performed at the following settings: initial denaturation at 95°C for 30s; 10 cycles of 95°C 30s, 49°C 60s, 68°C 100s; 24 cycles of 95°C 30s, T_a_ 60s, 68° at variable time, and a final extension at 68°C for 5min. T_a_ and variable times for each set of primers are reported in Table 1. A derivative of pKD46-Kan^R^ was used as 0.1229 is resistant to Amp^R^. Electroporation was used to construct *E. coli* 0.1229(pKD46) and MG1655(pKD46), using a Bio-Rad Gene Pulser II and following protocols recommended by the manufacturer. Colonies containing pKD46-Kan^R^ were selected on LB plates with kanamycin. Strains containing pKD46 were grown to an OD_600_ of 0.3, and L-arabinose was added to a final concentration of 0.2M. After incubation for 1 hr, cells were washed and electroporated with the pKD3-derived PCR product. Transformants were selected on LB plates with chloramphenicol. Knockouts were confirmed by PCR using primers ~200bp upstream and downstream of the gene, using standard PCR settings (initial denaturation at 95°C for 30s; 35 cycles of 95°C 30s, variable amplification temperature (T_a_) 60s, 68°C at variable time; and a final extension at 68°C for 5min). This strategy was followed for all the knockouts, including primers and temperatures specific for each gene (Table 1).

### Gibson cloning

The 2745-7950 bp region of p0.1229_3 was cloned into pBR322 (pBR322::p0.1229_3^2745-7950^), using Gibson cloning as previously described (69). Briefly, primer pairs were constructed containing 30 bp annealing to the pBR322 insert site and 30 bp that would anneal to p0.1229_3. DNA from 0.1229 and pBR322 was amplified at these sites using standard PCR settings, amplicons were cleaned up using a PCR purification kit (Qiagen, Germantown, MD) and subjected to assembly at 50°C using the Gibson cloning kit (New England Biosciences, Ipswich, MA). Assembled plasmids were propagated in DH5a competent cells (New England Biosciences, Ipswich, MA). Verification PCR was performed using primers 200 bp upstream and downstream of the insert site (Table 1) and confirmed using Sanger sequencing. Successful constructs were transformed into C600 electrocompetent cells. A similar process was used to clone *tolC* in pBAD18 (Kan^R^).

### Whole genome sequencing and bioinformatics

For the whole genome sequencing of 0.1229, genomic DNA was isolated using the Wizard Genomic DNA purification kit (Promega, Madison, WI). Whole genome sequencing was performed at the Penn State Genomics Core facility using the Illumina MiSeq platform. A PCR-free DNA kit was used for library preparation. The sequencing run produced 2 x 150 bp reads.

For the whole genome sequencing of 99.0750, 91.0593, and 90.2723, genomic DNA was isolated using Qiagen DNeasy Blood and Tissue Kit (Qiagen Inc., Germantown, MD). Whole genome sequencing was performed using the NexTera XT DNA library prep kit and run on an Illumina MiSeq platform. The sequencing run produced 2 x 250 bp reads.

After Illumina sequencing, Fastq files were checked using Fastqc v0.11.5 (70) and assembled using SPAdes v3.10 (71). SPAdes assemblies were subjected to the Quality Assessment Tool for Genome Assemblies v4.5 (QUAST) (72), and contig number, genome size, N50 and GC % were noted.

Strain 0.1229 was also sequenced at the Center for Food Safety and Nutrition, Food and Drug Administration using the Pacific Biosciences (PacBio) RS II sequencing platform, as previously reported (73). For library preparation, 10 μg genomic DNA was sheared to 20 kb fragments by g-tubes (Covaris, Inc., Woburn, MA, USA) according to the manufacturer’s instructions. The SMRTbell 20 kb template library was constructed using DNA Template Prep kit 1.0 (Pacific Biosciences, Menlo Park, CA, USA). BluePippin (Sage Science, Beverly, MA, USA) was used for size selection, and sequencing was performed using the P6/C4 chemistry on two single-molecule real-time (SMRT) cells with a 240 min collection protocol along with stage start. SMRT Analysis 2.3.0 was used for read analysis, and de novo assembly using the PacBio Hierarchical Genome Assembly Process (HGAP3.0) program. The assembly output from HGAP contained overlapping regions at the end which can be identified using dot plots in Gepard (74). The genome was checked manually for even sequencing coverage. Afterwards, the improved consensus sequence was uploaded in SMRT Analysis 2.3.0 to determine the final consensus and accuracy scores using Quiver consensus algorithm (75). The assembled genome was annotated using the NCBI’s Prokaryotic Genomes Automatic Annotation Pipeline (PGAAP) (76).

Plasmid sequences were visualized using Blast Ring Image Generator v0.95 (BRIG) (77). The Center for Genomic Epidemiology website was used for ResFinder v3.1.0 (90% identity, 60% length) (78), SerotypeFinder v2.0.1 (85% identity, 60% length) (79) and MLSTFinder v2.0.1 (80) using the Achtman multi-locus sequence typing (MLST) scheme (81). The Integrated Microbial Genomics & Microbiomes website of DOE’s Joint Genome Institute was utilized to BLAST the amino acid sequence of Hp1 against other genomes, matches that were between 36 and 68% identical from varying species were selected, then visualized using the gene neighborhoods function (82).

### Data Analysis

MS Excel (Microsoft Corporation, Albuquerque NM) was used to calculate the mean, standard deviation, and standard error; and GraphPad Prism 6 (GraphPad Software, San Diego CA) was used for generating figures. Error bars report standard error of the mean from at least three biological replicates.

### Data availability

Nucleotide and SRA files for the 0.1229 can be found on NCBI under Biosample SAMN08737532. SRA files for 99.0750 (SAMN11457477), 91.0593 (SAMN11457478), 90.2723 (SAMN11457479) can be found under their respective accession numbers.

## Acknowledgements

We thank Dr. Craig Stephens at Santa Clara University for providing strain SF-173, Dr. Roberto Kolter at Harvard Medical School for providing strain ZK1526, and Erin Nawrocki for manuscript proofreading. HF was supported by USDA National Needs Grant 2014-38420-21822. This work was supported by grant number 1 R21 AI130856-01A1 through the National Institute of Allergy and Infectious Diseases and the USDA National Institute of Food and Agriculture Federal Appropriations under project PEN04522 and accession number 0233376.

---

Fig. S1: W3110Δ*tolC* P*recA-gfp* was grown with 0.1229 and 0.1229 regional (A) or individual ORF (B) knockouts, or alone indicated by LB. One-way ANOVA was used and levels marked with an asterisk were significantly lower than 0.1229 (Dunnett’s test, p < 0.05).

Fig. S2: A portion of p0.1229_3 is compared to *K. pneumoniae* TR152 (SAMEA3729690), *S. sonnei* 143778 (SAMEA2057991), *E. coli* HUSEC41 (PRJEA73977), *E. coli* HVH206 (SAMN01885845). * indicates one amino acid (aa) difference in that ORF. # indicates seven aa differences in that ORF. The grey shaded region is >99.6% nucleotide identical between all strains.

Fig. S3: Hp1 protein was compared to genomes on Integrated Microbial Genomes & Microbiomes of DOE’s Joint Genome Institute. Using BLASTp, isolates were compared, sorted by BIT score, and then one strain from the top ten species were selected. % identity ranged from 32 to 68%. Hp1 homologs are colored in red. 8/10 have ABC transporters adjacent to Hp1, colored in light blue or maroon. 9/10 have an annotated region similar to Cupin, colored in pale yellow, typically adjacent to the ABC transporter. DUF2164 is found in five strains, one in the reverse direction than Hp1. The *Burkholderia cepacia* strain encodes L-arabinose system upstream of Hp1. The bracket indicates groupings of Hp1, ABC and Cupin.

Fig. S4: W3110Δ*tolC* P*recA-gfp* was grown with 0.1229 or 101 human fecal isolates from the *E. coli* Reference Center at Penn State. As a control, W3110 P*recA-GFP* was grown by itself indicated by LB. One-way ANOVA was used and levels marked with an asterisk were significantly higher than LB (Dunnett’s test, p < 0.05).

